# Mind of a dauer: Comparative connectomics reveals developmental plasticity

**DOI:** 10.1101/2023.03.23.533915

**Authors:** Hyunsoo Yim, Daniel T. Choe, J. Alexander Bae, Hae-Mook Kang, Ken C.Q. Nguyen, Myung-kyu Choi, Soungyub Ahn, Sang-kyu Bahn, Heeseung Yang, David H. Hall, Jinseop S. Kim, Junho Lee

## Abstract

A fundamental question in neurodevelopmental biology is how flexibly the nervous system changes during development. To address this, we reconstructed the complete connectome of dauer, an alternative developmental stage of nematodes with distinct behavioral characteristics, by volumetric reconstruction and automated synapse detection using deep learning. With the basic architecture of the nervous system preserved, structural changes in neurons, large or small, were closely associated with connectivity changes, which in turn evoked dauer-specific behaviors such as nictation. Graph theoretical analyses revealed significant dauer-specific rewiring of sensory neuron connectivity and increased clustering within motor neurons in the dauer connectome. We suggest that the nervous system in the nematode, probably animals in general, has evolved to respond to harsh environments by reversibly developing a quantitatively and qualitatively differentiated connectome.

## Introduction

A fundamental question in neurobiology is how the nervous system governs behavior (Emmons, 2015). Specifically, researchers seek to understand how an animal’s nervous system assimilates diverse sensory information from its external surroundings to make adaptive decisions for survival. Elucidating and analyzing connectomes of animals is one way to address this question. A more specific question in neurobiology in terms of development concerns the capacity of an animal to dynamically shape its nervous system throughout its development. In other words, how extensively does an animal modify its neuronal connections in response to physiological and external cues at different developmental stages? Comparative connectomics provides excellent opportunities to answer this question of developmental plasticity.

Since Sydney Brenner initiated his ambitious research on the connectome, the nematode *C. elegans* has been at the center of connectomics studies (Emmons, 2015; White et al., 1986). Connectomics studies using *C. elegans* have been successfully extended to comparative studies, comparing the adult nervous systems of both sexes (Cook et al., 2019; Jarrell et al., 2012) and investigating the neural development along the series of life stages (Witvliet et al., 2021). Along the same lines of research directions, but more focused on developmental plasticity, the study of the connectome of dauer, an alternative developmental stage, is particularly intriguing. Under unfavorable conditions, *C. elegans* enters the diapause state of dauer, and systematic anatomical changes occur to many tissues, acquiring physiological robustness and displaying various unique behavioral repertoires (Figure 1A; Altun and Hall, 2009; Cassada and Russell, 1975; Gaglia and Kenyon, 2009; Golden and Riddle, 1984a; b; Hallem et al., 2011). The remodeling of the connectome underlying these behavioral changes in dauer could provide insights on both the neural substrates of behaviors and the principles of the plasticity of animal development (Albert and Riddle, 1983; Britz et al., 2021).

**Figure 1.**
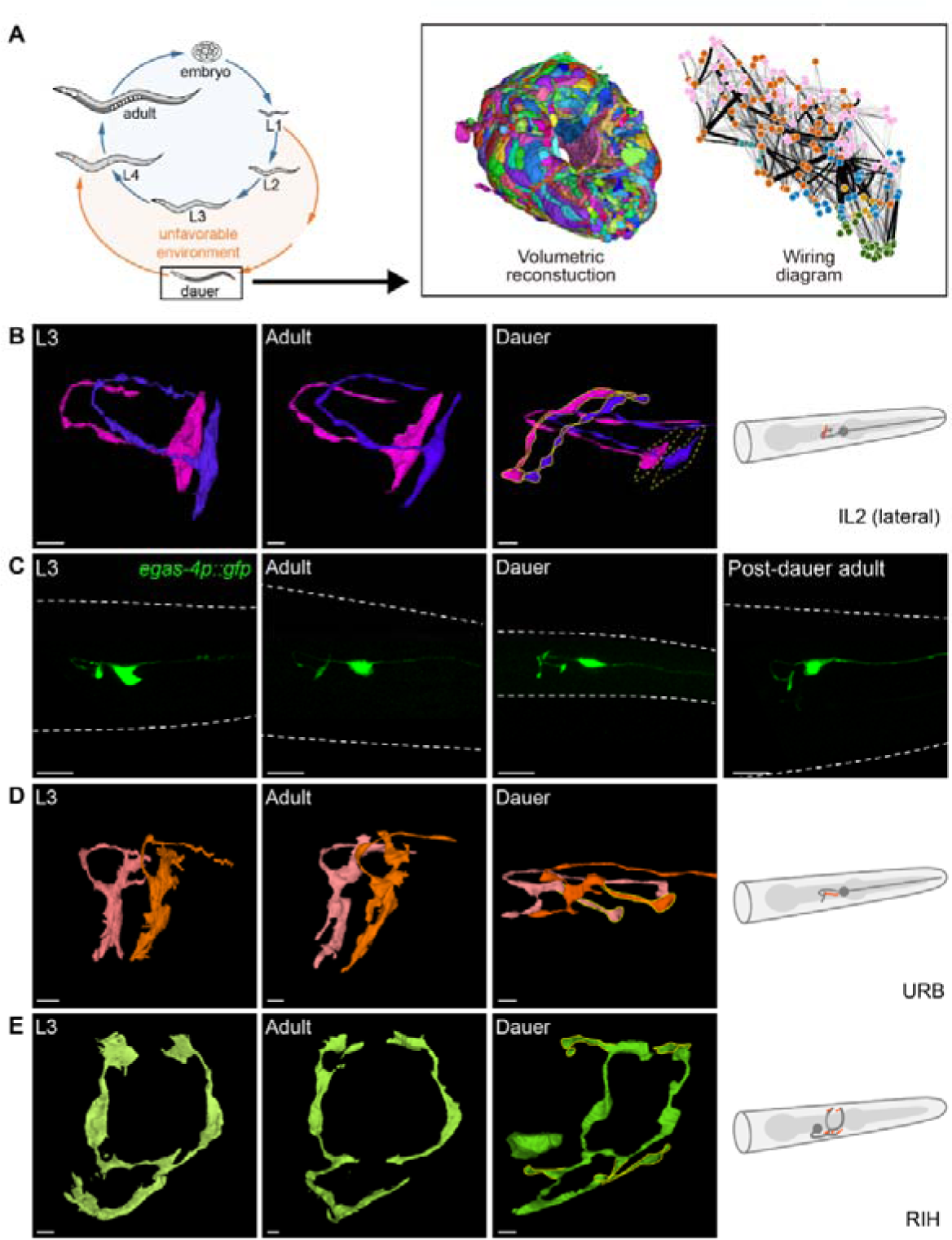
Volumetric reconstruction of the dauer nerve ring. (A) Volumetric reconstruction and wiring diagram (pink: sensory neuron, orange: interneuron, blue: motor neuron, green: body wall muscle) of dauer nerve ring. Volumetric reconstruction is colored to distinguish individual neurons. Arrangement of nodes in the wiring diagram is adopted from (Witvliet *et al*., 2021). (B) Volumetric reconstruction of IL2 lateral neurons in L3, adult, dauer (from left) and schematic view of neuron in dauer stage (right). New branches emerge from the axon (solid yellow), and the terminal swelling shrinks (dashed yellow). (C) Fluorescence images of IL2 lateral neurons in L3, adult, dauer, and post-dauer adult (from left). IL2 lateral neurons were marked with GFP using *egas-4* promoter. (D) Same as (B) for URB neurons. New branches emerge from terminal swelling (solid yellow). (E) Same as (B) for RIH neurons. Small branches emerge from various regions (solid yellow). (B, D, and E) Scale bar: 1 μm. (C) Scale bar: 10 μm. See also Figures S1 and S2.

In this study, we report the complete connectome of a hermaphrodite dauer nerve ring (Figure 1A), in comparison with the publicly available 10 connectome datasets consisting of L1∼L4 and adult stages from previous studies (Figure S1A; (Brittin et al., 2021; Cook *et al*., 2019; Cook et al., 2023; Witvliet *et al*., 2021). We show that local and modest, but significant, morphological changes of dauer neuronal arbors accompany synaptic changes, while most synapses remain intact through development. We also report the results of the graph theoretical analyses of the dauer connectome for the purpose of identifying the structural changes in the network topology that affect the dynamics of the overall neural network. Our work implies that substantial, reversible changes in behavior can be accomplished within a static set of neurons, through plastic alterations among their local synaptic connections.

## Results

### Volumetric reconstruction of dauer nerve ring reveals morphological changes in neurons

A *C. elegans* dauer nerve ring sample was sectioned and imaged at every 50 nm, resulting in 364 serial TEM images. The dense volumetric reconstruction (Figures 1A and S1B; STAR Methods) yielded the processes of 181 neurons in the nerve ring, whose number is only modestly different from those of the other late developmental stages (Figure S1A). After the volumetric reconstruction and cell identification, we visually inspected the neural processes using the 3D renderings (Table S1). We noticed the changes in dauer with varying magnitudes and forms, the most prominent of which were categorized into Branching (B), small Protrusion (P), Retraction (R), and Shrinkage (S) of the neurites and terminal swellings. Unlike the minor anatomical changes except for growth in size observed across other development stages (Witvliet *et al*., 2021), the changes in dauer were often more dramatic as shown in Figure S2.

The IL2 neurons are a family of six sensory neurons, critical to the dauer-specific phoretic behavior called nictation; two of these are lateral cells and the rest are quadrant cells (dorsal and ventral) (Lee et al., 2011; White *et al*., 1986). Dauer IL2 lateral neurons add an extra branch that emerges perpendicularly from the apex of an axon bend (Figures 1B and 1C). Previous literature reported that while IL2 quadrant neurons undergo significant remodeling in their processes during dauer development, IL2 lateral neurons change less so (Schroeder et al., 2013). Thanks to the finer resolution of TEM, we discovered that remodeling occurs in IL2 lateral neurons as well. As a validation, we employed fluorescence microscopy to observe the IL2 lateral neurons in different stages including a post-dauer adult (Figure 1C), confirming that the extra branches of IL2 lateral neurons appear only in dauer. In addition, the terminal swellings of IL2 lateral neurons were shrunken in dauer, losing contact with the IL2 dorsal neurons (Figure 1B). The URB neurons (Figure 1D) and RIH neurons (Figure 1E) also show distinctive structural changes in dauer.

### Dauer connectome exhibits similarities and differences with other stages

In order to see how synaptic connections change according to the morphological changes, we analyzed the full nerve ring connectome of the dauer. We annotated the chemical synapses in the image stack, yielding a complete connectivity map between the reconstructed neurons (Figure 1A) (Altun and Hall, 2009; Witvliet *et al*., 2021). A convolutional neural network (CNN) was trained to predict the locations of active zones (STAR Methods) and the predicted active zones were segmented to distinguish each chemical synapse. The postsynaptic partners of an active zone, together forming synapses, were assigned by simulating neurotransmitter diffusion to the closest neighboring axons (STAR Methods; (Witvliet *et al*., 2021).To ensure high accuracy, the detected synapses were proofread by human experts (Figures S1B and S1C; STAR Methods).

All together, 2,813 active zones and 6,371 synapses were found in the dauer nerve ring (Figures S3A and S3B). Each active zone pairs with 2.26 postsynaptic partners on average (SD=0.93; Figure S3C). 19% of the active zones form monadic and 81% polyadic synapses. The 6,371 synapses form 2,200 connections, meaning that a pair of pre- and postsynaptic neurons are connected by 2.90 synapses on average (Figure S3D). As the size of the active zone could be a proxy for the functional strength of the synapse, the sum of active zone sizes is regarded as the connection weight or strength between two neurons. In summary, the dauer connectome is represented by a directed weighted graph with 181 nodes of neurons and 1,995 neuronal connections, or in total 2,200 connections including the neuromuscular junctions (Figure 2A, Table S2).

**Figure 2.**
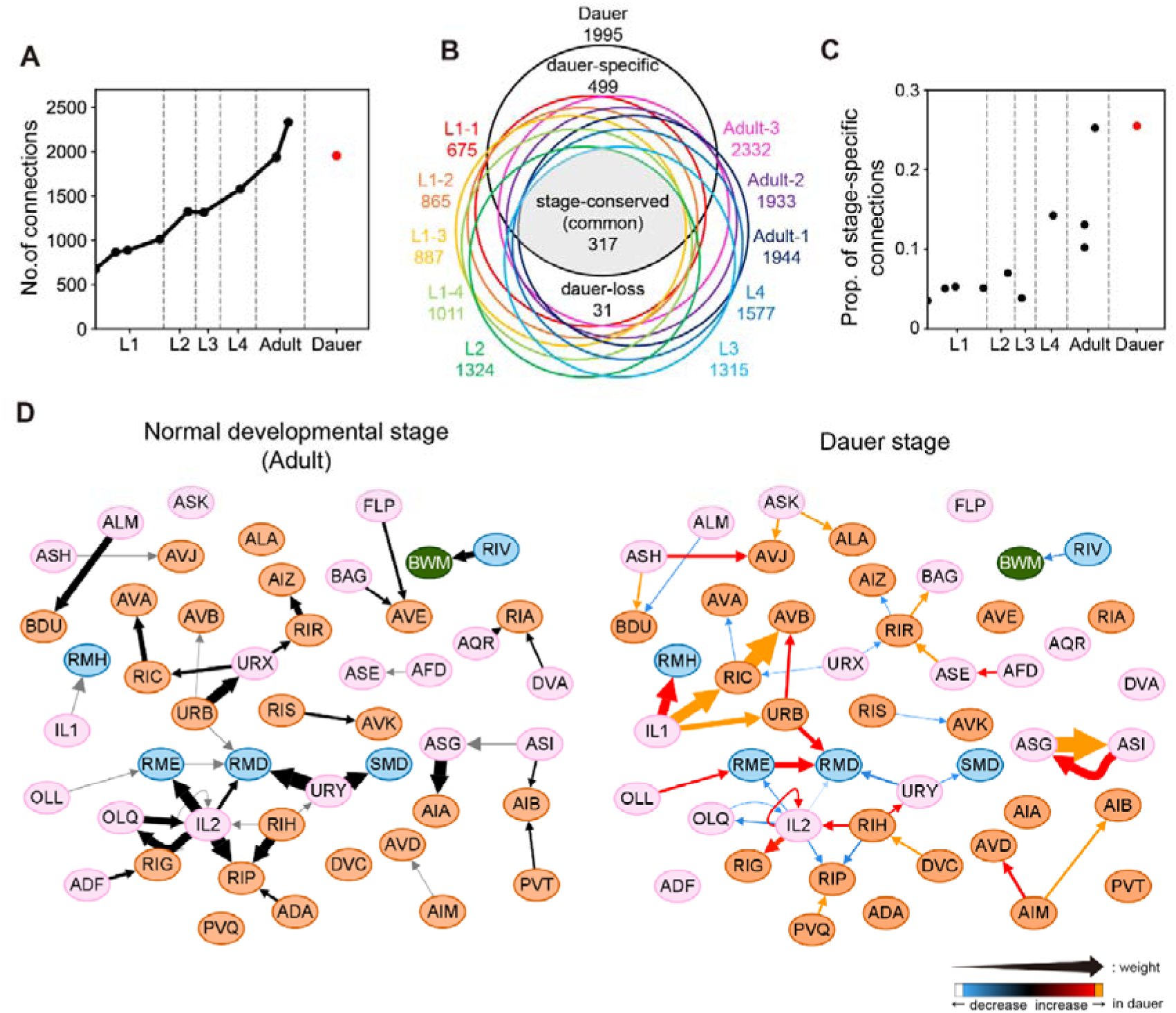
Completion of dauer connectome. (A) Total number of connections between neurons in different developmental stages. Total number of connections in dauer is similar to that of an adult. (B) Venn diagram of connections between neurons. 499 connections are connections that only exist in dauer. Area of each region is not scaled according to the number of connections for visual purposes. (C) Proportion of stage-specific connections within each dataset. Over a quarter of connections are dauer-specific connections in dauer, which are considerably higher than other stages. (D) Wiring diagram that includes representative changes in dauer connections (right) and wiring diagram of adult stage (left) for the same set of neurons (pink: sensory neuron, orange: interneuron, blue: motor neuron, green: body wall muscle). The width of arrow indicates the relative connection weight and colors indicate changes in dauer relative to adult stage. Dauer-specific connections are represented with orange arrows. Connections which connection weight increased and decreased in dauer are represented with red and blue arrows respectively. See also Figure S3.

We could now compare the connectivities of dauer with other stages from earlier studies (Figure S1A; (Brittin *et al*., 2021; Cook *et al*., 2019; Cook *et al*., 2023; White *et al*., 1986; Witvliet *et al*., 2021). We found that the total number of connections in the dauer nerve ring is similar to that of the adult nerve ring, thus deviating from the trend of linear gains with increasing larval ages (Figure 2A; (Witvliet *et al*., 2021). Comparisons across all developmental stages, including dauer, show 317 connections in common, which we call the “stage-conserved” connections (Figure 2B). Another 499 connections exist only in the dauer, which we call the “dauer-specific” connections (Figure 2B; Table S3). On the other hand, 31 connections exist only in other stages but not in dauer and we call these “dauer-loss” connections (Figure 2C; Table S3). Interestingly, dauer has over 25% of stage-specific connections, in contrast to the cases of other stages which have roughly 9% of stage-specific connections on average (Figures 2C and S3E). There are also many connections that are shared with other stages but show different connection weights in dauer (Figure 2D). Wiring diagram differences of all individual neurons are summarized in Supplementary item 1.

### Morphological changes and connectional changes are mutually associated

We examined whether the major morphological changes listed in Figure 1 and Figure S2 were associated with the formation of new synapses (Figures S2 and S3). We found that all the morphological changes described in Figure S2 were accompanied with the formation of new synaptic inputs and outputs. For example, the IL2s with extra arbors established new IL2➔AUA and DVC➔IL2 synapses (Figures S4A and S4B). The dauer URB neuron received new synaptic input from IL1 on its new arbor on the dorsal side (Figure S4C), which forms one of the largest dauer-specific connections (Figure S4D). RIH neurons grew 5 new branches, where a multitude of new synaptic inputs and outputs were made with various partners, among which the DVC➔RIH synapse was notable (Figures S4E and S4F). Likewise, all the morphological changes described in Figure S2 were accompanied with the formation of new synaptic inputs and outputs.

Since some synaptic changes could involve less obvious morphological changes, we then inspected whether the connections with significant changes in their weights are associated with any morphological changes. We found that, for most connection changes described in Figure 2D, thickening of neural processes or broadening of local swellings contributed to the changes in contact area and resulted in the establishment of new synapses. The connection with the largest weight in all dauer connections was ASG➔ASI connection, both of which are amphid sensory neurons known to be involved in the dauer-entry decision (Bargmann and Horvitz, 1991). The ASI➔ASG connection weight was also dramatically increased in dauer (Figure 3C, Table S3). ASG and ASI did not form any synapse with each other in any reproductive stage although they did form several local contacts (Figure 3A). At the corresponding location in the dauer, a new swelling developed in ASI while ASG tightly adhered to the swelling, producing a larger contact with new large synapse (Figures 3A and 3C). The total ASI➔ASG contact area in the dauer increased by a factor of 2.54 compared to adult (STAR Methods; Table S4). ASG output connections exhibited large weight loss in dauer. The ASG➔AIA connection standed out among the 31 dauer-loss connections, as it is a strong connection in other developmental stages which has totally disappeared in dauer (Figure 3B; Table S3). The size of the local swelling in AIA decreased and the contact area between AIA and ASG also decreased. Thus, the total contact area between ASG and AIA was reduced by a factor of 0.33 in dauer compared to adult (Table S4). We have observed a similar correlation between connectivity and contact in IL1 ➔ RIC connection (Figures 3D and 3F) and RIC ➔ AVA and AVB connections (Figures 3E and 3F). In dauer, the total contact area between IL1 and RIC increased by a factor of 8.14 representing strengthened IL1➔RIC connection. Moreover, the total contact area of RIC and AVA decreased and that of RIC and AVB increased by a factor of 0.5 and 6.16, respectively, which were consistent with connectivity changes.

**Figure 3.**
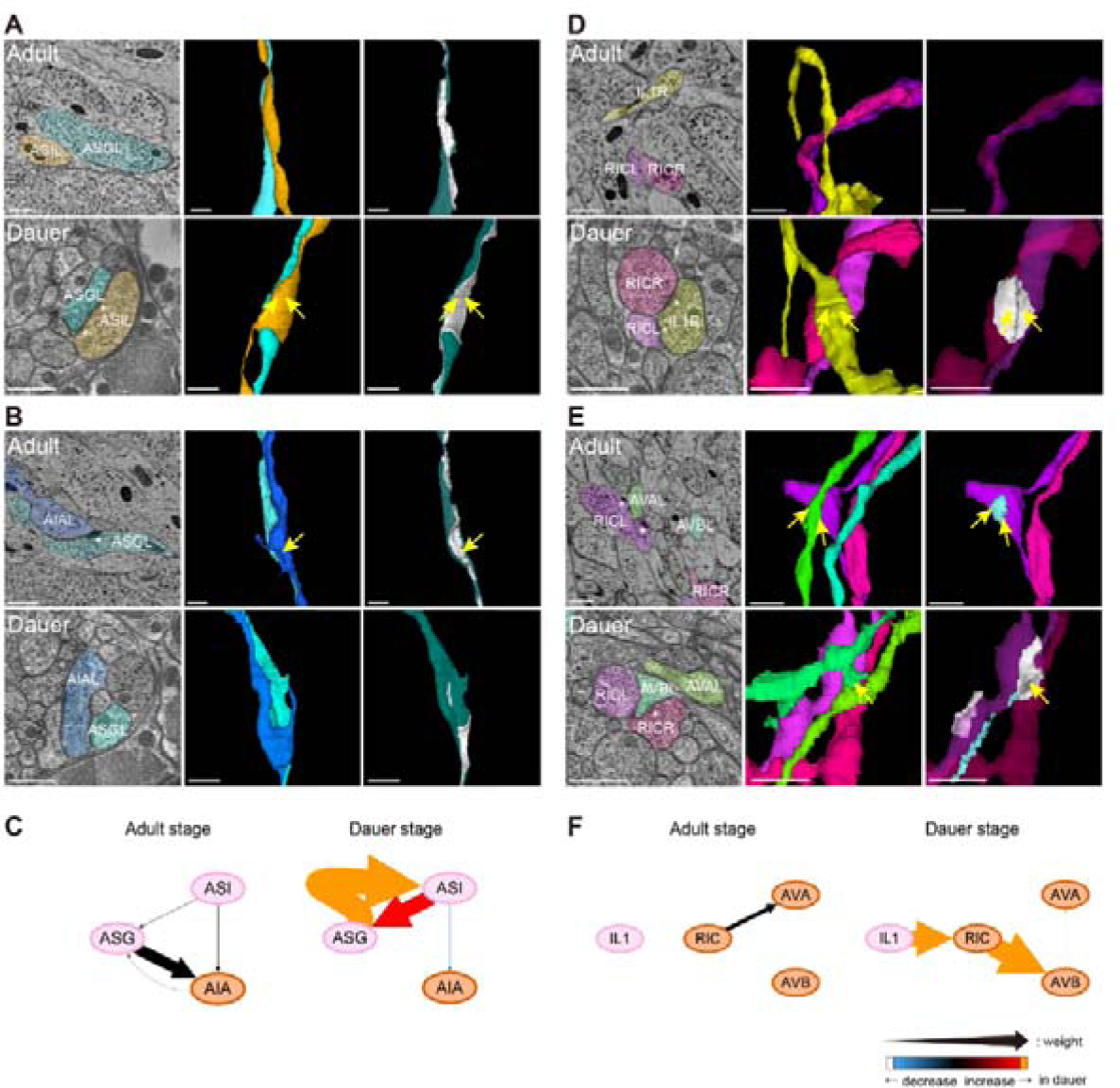
Morphological changes and connectional changes are mutually associated. (A) EM cross-section view (left) and 3D volumetric reconstructions of presynaptic and postsynaptic neurons (middle) at the location of ASG-ASI contact where synapse exists (right, contact area marked as white) in adult (top row) and dauer (bottom row). Contact area between ASG and ASI was significantly larger in dauer resulting in dauer-specific synapses (yellow arrows). Note that EM cross-section views of two stages may look different as two datasets have been sectioned in different orientations. (B) Same with (A) for ASG-AIA contact. ASG-AIA contact area is significantly reduced in dauer while ASG-AIA contact area is distinctive resulting in larger synapse (yellow arrow). (C) Wiring diagram of neurons shown in (A) and (B) in adutl stage (left) and dauer stage (right). Width and color of the arrow follows the same rule from Figure 2D. Larger contact between ASG and ASI results in strong reciprocal connections and smaller contact between ASG and AIA results in no connection in dauer. (D) Same with (A) for RIC-IL1 contact. IL1-RIC contact exists in the dauer stage while they do not make contact in the adult stage resulting in dauer-specific synapses (yellow arrows). (E) Same with (A) for RIC-AVA (light blue) and RIC-AVB (white) contacts. Area of RIC-AVA contact decreased in the dauer stage, leading to lack of synapses in the dauer stage while RIC-AVA synapses (yellow arrows) exist in the adult stage. Area of RIC-AVB contact increased in the dauer stage resulting in dauer-specific synapses (yellow arrow). (F) Wiring diagram of neurons shown in (D) and (E) in adult stage (left) and dauer stage (right). Contact between RIC-IL1 and RIC-AVB results in new connections and lack of contact between RIC-AVA results in no connection in dauer. (A, B, D, E) Scale bars: 500 nm (EM images) and 1 μm (3D views), asterisk: active zone, yellow arrow: stage-specific synapse. See also Figure S4.

### Connectivity changes are correlated with unique behavior in dauer

Next, we asked whether the connectivity changes in dauer are associated with the unique behavior of dauer. We first focused on the IL2 neurons, which are known to play an essential role in nictation (Lee *et al*., 2011). We found that IL2 quadrant neurons established strong connection to RIG neurons in dauer compared to other stages (Figure 4A). In dauer, each IL2 process forms a new swelling, forming new contact and large synapse with RIG neurons (Figure 4B). Therefore, we hypothesized that RIG could be potential neurons involved in the neural circuit regulating nictation, serving as downstream neurons of IL2 neurons. To test this, we performed behavioral tests with dauers bearing mutations in two genes, *cha-1* and *daf-10*, which are known to show poor nictation (Lee *et al*., 2011). The nictation ratio of *cha-1* and *daf-10* mutant dauers was significantly increased by optogenetic activation of RIG (Figures 4C and 4D), suggesting that RIG activation is sufficient for the nictation of dauer even in the absence of ciliated, functional IL2 neurons and that RIG neurons act downstream of IL2 in nictation. Consequently, strengthened anatomical IL2➔RIG connection in the dauer stage is believed to have a functional role leading to behavior.

**Figure 4.**
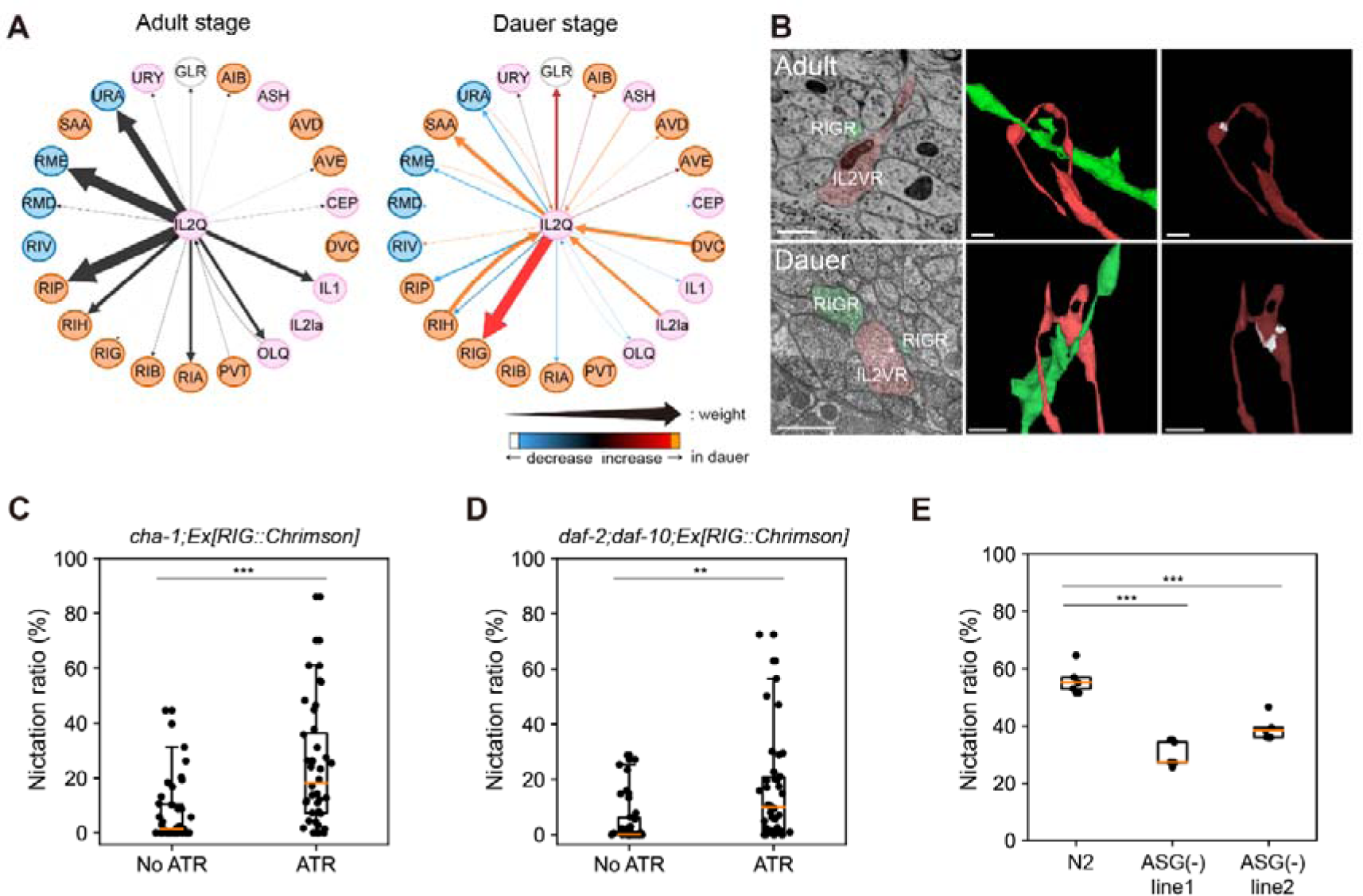
Connectivity changes in dauer neurons are correlated with unique behavior in dauer. (A) Wiring diagrams showing connections of IL2 quadrant neurons. Connection from IL2 to RIG shows significant increase in strength in the dauer stage. Width and color of the arrow follows the same rule from Figure 2D. (B) EM cross-section view (left) and 3D volumetric reconstructions of presynaptic and postsynaptic neurons (middle) at the location of IL2Q-RIG contact (right, contact area marked as white) in adult (top row) and dauer (bottom row). (C) RIG-activated *cha-1* mutants exhibit significantly higher nictation ratio than*cha-1* mutants (individual nictation test; *n*=40; unpaired t-test; *p*<0.0001). (D) RIG-activated *daf-2*;*daf-10* mutants exhibit significantly higher nictation ratio than*daf-2*;*daf-10* mutants (individual nictation test; *n*=38; unpaired t-test; *p*=0.0016). (E) ASG ablated lines show significantly lower nictation ratio than N2 (population nictation test; 5 trials, *n*>60 worms per experiment; unpaired t-test;*p*<0.0001). (B) Scale bars: 500 nm (EM images) and 1 μm (3D views), asterisk: active zone. (C-E) Orange line: median, box: interquartile range, whiskers: 5th and 95th percentiles.

Another candidate neurons for dauer-specific physiology and behavior were the ASG neurons as the connections of ASG showed the most dramatic change in strength (Figure 3A). We found that ASG-ablated worms exhibit deficit in nictation behavior, suggesting that ASG could also be associated with nictation (Figure 4E). In summary, our investigation of individual neuronal changes within the connectome revealed that the changes in morphology of neurons in dauer are closely related to connectivity changes, which could explain the stage-specific behavioral changes.

### The networks of dauer and adult share common structural properties

Next, we compared the entire dauer connectome with connectomes of other stages systematically using graph theoretical analyses, seeking global characteristics of the dauer connectome. We aforementioned the dauer connectome has a similar number of connections with adult connectome (Figure 2A). Consistently, the average in- and out-degrees of neurons both exhibit increase across developmental stages and the dauer connectome shows similar values with adult connectomes (Figure 5A). Furthermore, when we look at the degree distributions of each stage, the distributions decay roughly exponentially as revealed by linear decrease in the semi-log plot (Figure 5B). The exponential decay implies that the network mainly consists of peripheral neurons and only a few small hub neurons such as RIH, AVB, AIB, and RIA. In addition, the distributions are shifted toward larger values and tails tend to become longer for dauer and adult-stage networks (Figure 5B), suggesting hub neurons tend to have more connections in the dauer and adult stages. As dauer and adult stages consist of more connections, it leads to shorter mean path length (Figure 5C) and higher clustering coefficient (CC), which measures the probability of the pair of neurons that are connected to a common neuron being connected to each other (Figure 5D). Overall, the dauer connectome shares common global network structures with adult connectomes.

**Figure 5.**
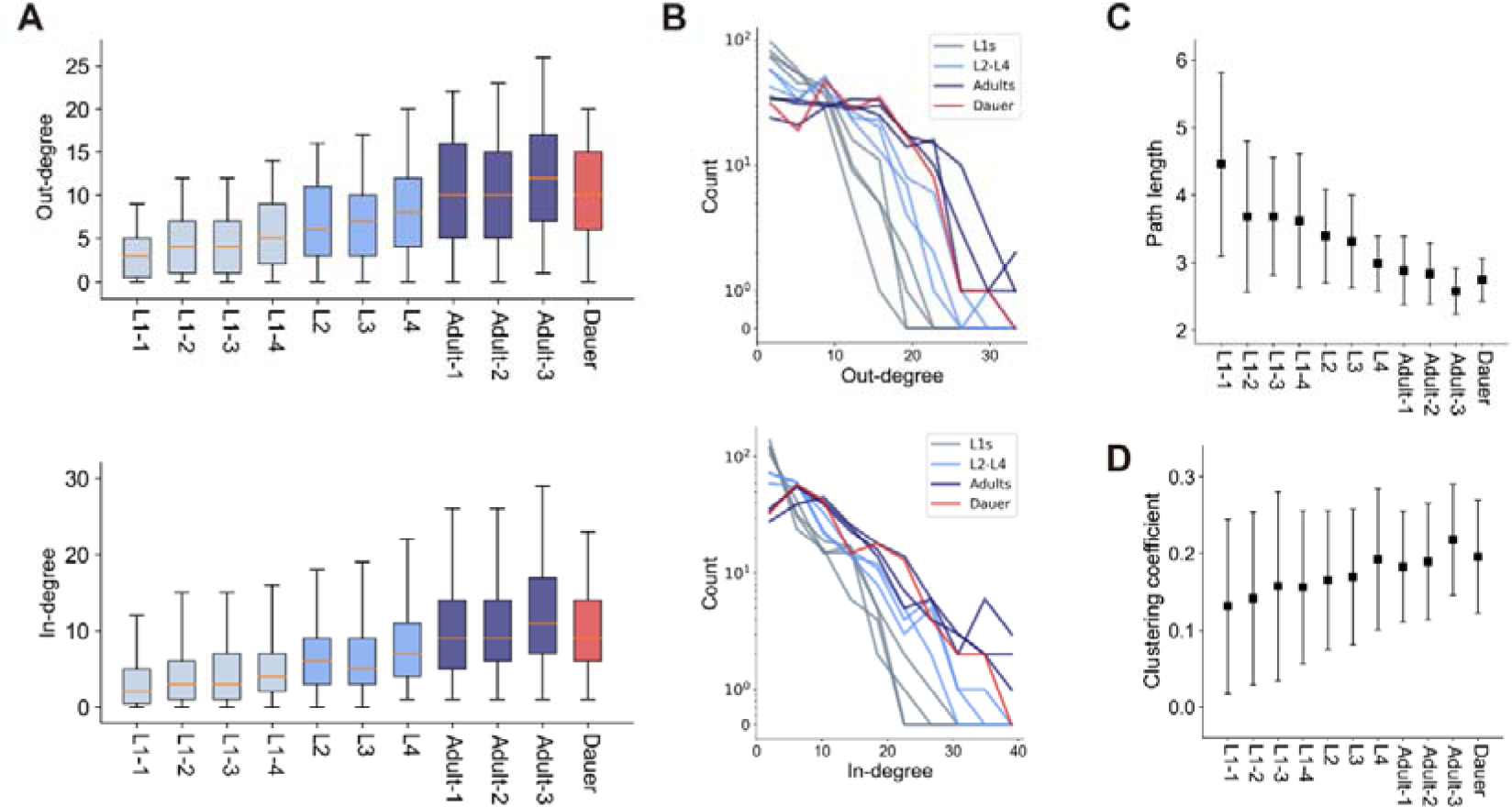
The networks of dauer and adult stages share common structural properties. (A) Out- (top) and in-degrees (bottom) of neurons (*n*=187; orange line: median, box: interquartile range, whiskers: 5th and 95th percentiles) across development. Boxes are colored by developmental stages (gray: L1s, light blue: L2 to L4, dark blue: adults, red: dauer). (B) Out- (top) and in-degree (bottom) distributions of different developmental stages. Line color follows the same rule in (A). (C) Mean path lengths of neurons (*n*=140, 142, 143, 145, 153, 152, 168, 159, 160, 177, 164; mean±SD) across development. (D) Clustering coefficients of neurons (*n*=155, 161, 158, 164, 172, 174, 178, 180, 180, 179, 179; mean±SD) across development.

### Dauer-specific connections contribute to increased clustering within motor neurons

While the dauer and adult stages share common features in primary network properties, we further explored whether there are differences between dauer and adult stages in subnetwork properties. We first divided the connections into 9 types according to the types of pre- and postsynaptic neurons (i.e. sensory, inter-, motor neurons) and measured the out-degree of neurons in each connection type (Figure 6A; (Cook *et al*., 2019). The out-degree of each connection type did not differ in most connection types except for M➔M connections, where dauer shows significantly greater out-degree. This indicates notable proportion of dauer-specific connections (Figure 2C) contribute to addition of connections within motor neurons. Thus, we inspected how these additional connections affect the network characteristic.

**Figure 6.**
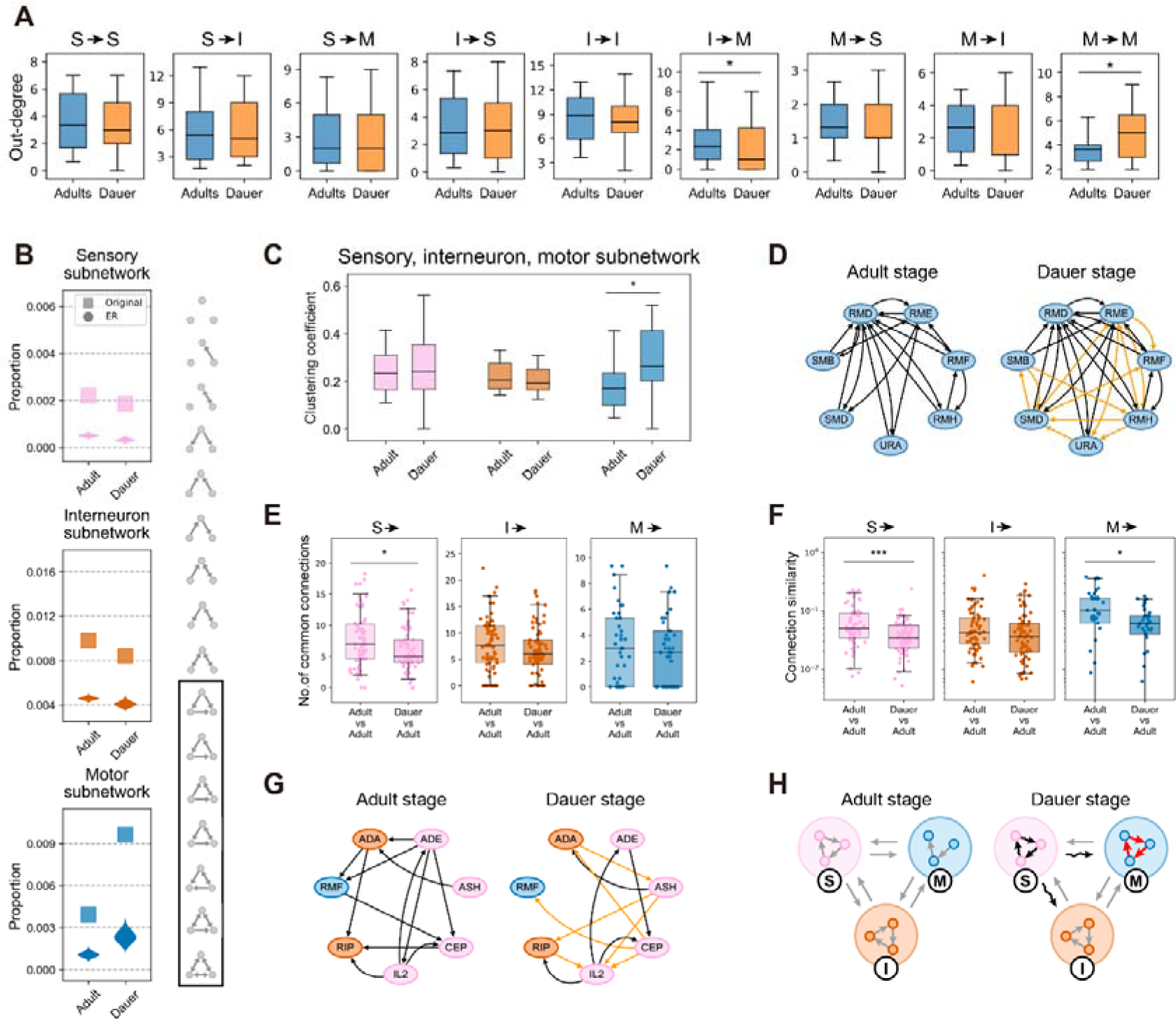
Dauer connectome exhibits rewired sensory connectivity and increased clustering within motor neurons. (A) Out-degrees of neurons for different connection types (Wilcoxon rank-sum test;*n*=156, \**p*<0.05) in adults (blue) and dauer (orange). Neurons without any connection in any of the datasets were excluded. (B) Observed proportion of triangular motifs (motifs in black rectangle; square) in sensory (pink), interneuron (orange), motor (blue) subnetworks in adults and dauer and their expected distributions in corresponding Erdős‒Rényi random networks (violin plot). (C) Clustering coefficients of neurons in sensory (pink), interneuron (orange), motor (blue) subnetworks in adults and dauer (*n*=63 (S, adult), 64 (S, dauer), 72 (I, adult), 73 (I, dauer), 31 (M, adult), 31 (M, dauer); Wilcoxon rank-sum test; \**p*<0.05, \*\**p*<0.01). Neurons with less than 2 connections in the subnetwork were excluded. (D) Wiring diagrams of representative motor neurons in adult and dauer stages. Dauer-specific connections (yellow) add more connections in the motor subnetwork, leading to higher local clustering. (E) Number of common output connections from sensory (pink), inter- (orange), motor (blue) neurons between adults (left) and that between dauer and adults (right; Wilcoxon rank-sum test; \**p*<0.05). Neurons without any connection in both datasets of the pair were excluded. (F) Connection similarity of output connections from sensory (pink), inter- (orange), motor (blue) neurons between adults (left) and that between dauer and adults (right; Wilcoxon rank-sum test; \**p*<0.05, \*\*\**p*<0.001). Neurons without any connection in both datasets of the pair were excluded. (G) Wiring diagrams showing example rewired sensoryneurons in adult and dauer stages. Dauer-specific connections (yellow) add new connections while losing connections that exist in adults, leading to rewired sensory connectivity. (H) Summary diagram illustrating differences between dauer and adult networks. (A, C, E, F) Black line: median, box: interquartile range, whiskers: 5th and 95th percentiles. (A, B, C) Values in adult are computed by averaging over three adult datasets. (E and F) Values are computed by averaging over all pairs of datasets. See also Figures S5 and S6.

To characterize the M➔M connections, we considered three types of subnetworks, each consisting of connections within a single type of neurons. The motor subnetwork was analyzed in comparison to sensory and interneuron subnetworks. We first performed three-node motif analysis in each subnetwork and discovered that the proportions of the triangular motifs, where all three neurons in a motif are connected to one another, were higher compared to the rest of the motifs (Figure 6B, STAR Methods). As a reference, the proportions of triangular-motifs were also measured in the equivalent Erdős‒Rényi (ER) random networks with the same number of nodes and average degrees. The motor subnetwork had highest proportion in dauer, indicating the connections of motor neurons of dauer are wired to make more triangular motifs compared to motor neurons of adult (Figure 6B). Motor neurons had the highest proportion compared to other types of neurons in dauer while this trend was not found in adults (Figure 6B). This is not due to the higher connection probability, since the equivalent ER random network could not reproduce the similar difference in triangular-motif proportion between dauer and adults in motor subnetwork (Figure 6B). This evidence highlights that dauer-specific connections between dauer motor neurons are programmed to make an exceptionally large number of triangular motifs.

As a validation, we computed the CCs of neurons in the three subnetworks and the neurons in the dauer motor subnetwork exhibited significantly larger CC than adult motor subnetworks while there were no differences in sensory and interneuron subnetworks (Figure 6C). This further supports the conclusion that dauer motor neurons are specifically connected for local clustering. The subnetwork diagrams of a few major motor neurons for adult and dauer exemplify that the dauer-specific connections enhance the proportion of triangular motifs and the clustering among motor neurons in dauer (Figure 6D).

### Dauer-specific rewiring of sensory neuron connectivity

Unlike motor neurons, other types of neurons did not show significant differences in out-degree (Figures 6A and S5A). Considering the high proportion of dauer-specific connections (Figure 2C), we hypothesized the connection target distribution could differ for sensory neurons and interneurons leading to rewiring in connectivity.

We first analyzed the number of common connections between the datasets. We found that the number of common connections in sensory neuron output connections between dauer and adult datasets is significantly smaller than the number of common connections between adult datasets (Figure 6E, S5B-S5D). On the other hand, the number of common connections in inter- and motor neuron output connections between dauer and adult datasets did not exhibit differences with the number of common connections between adult datasets (Figures 6E, S5B-S5D). This means the smaller number of common connections from sensory neurons is a stage-specific phenomenon in contrast to inter- and motor neurons.

The fact that they have fewer common connections is not absolute evidence for the rewiring in output connectivity of sensory neurons in dauers, because there could be neurons that are not rewired but only have few connections all shared between datasets. Therefore, to supplement our argument of rewiring, we introduced “connection similarity (CS)”, which measures the similarity of connection targets of neurons between two stages (STAR Methods). We found that CS between dauer and adult stages are significantly lower than CS among adult stages in sensory neurons while this is not the case for interneurons (Figure 6F and S5E). This result implies that output connections of sensory neurons are dauer-specifically rewired compared to adults while both have similar number of connections. This result is consistent with results regarding the number of common connections (Figures 6A, 6F, S5B, S5C, and S5E). For interneurons, CSs exhibited a similar level for every pair of datasets indicating it is due to variability between datasets rather than dauer-specific phenomenon (Figures 6A, 6F, and S5E). For motor neurons, CSs of dauer and adults were lower than CSs between adults unlike the results regarding the number of common connections. This provides additional evidence that dauer-specific connections contribute to additional connections on top of common connections for motor neurons in dauer (Figures 6A and 6F).

In conclusion for the graph theoretical analyses, there exists increased clustering among motor neurons and significant rewiring in the connectivity of sensory neurons in the dauer connectome, creating differences with adult connectomes (Figures 6D, 6G, 6H).

## Discussion

Complete stage-wise comparison of neural circuits at the single synapse level is becoming feasible as EM connectome datasets in all developmental stages have been published (Brittin *et al*., 2021; Cook *et al*., 2019; Cook *et al*., 2023; White *et al*., 1986; Witvliet *et al*., 2021) except for the alternative stage, dauer. As the last piece of the puzzle, we have reconstructed the first dauer nerve ring connectome from EM images, consisting of dense volumetric reconstructions of both neuronal and muscle cells, and their connectivity acquired using deep learning. Using the publicly available resource, we have found that various morphological changes of neurons occur at the dauer stage and these morphological changes are related to connectivity changes. Also, we have shown that dauer-specific connections are associated with the unique survival behavior, nictation. Lastly, we found that the neural network of dauer exhibits rewired sensory connections and increased clustering among motor neurons.

Morphological changes associated with synaptic changes must not be a mere coincidence but must be an event with concrete physiological advantages, because morphological remodeling is a highly energy-consuming process (Chi et al., 2022; Steiner, 2019). The local morphological changes are permissive for connectional changes, and these changes can lead to valuable behavioral changes, even without changes in the number of neurons involved. The changes in morphologies and connections can be identified by comparative connectomics. The original question that Brenner posed, how the nervous system governs the behavior, can now be recast as the relationship between dauer-specific connections and unique behaviors of the dauer, which can be further explored by future experimental studies. One example is CO_2_ sensing in different developmental stages. Consistent with a previous report that AIY is not required for CO_2_ response in dauers unlike adults (Banerjee et al., 2023), we easily identified that the connection from BAG (Hallem *et al*., 2011), which is responsible for CO_2_ detection, to AIY does not exist in the dauer connectome while it exists in the adult. Now, it is possible to focus on additional stage-specific connections and check how the neuronal functions differ in terms of CO_2_ sensing using optogenetics and calcium imaging like the IL2-RIG experiments we have conducted. Another example is the connection changed from RIC➔AVA in other larval stages to RIC➔AVB in the dauer. Proper analyses and experiments will reveal the significance of this shift in dauer-specific motility. As these examples demonstrate, we believe that our new dauer connectome could lead to new discoveries of neural mechanisms for dauer-specific functions.

Along with other developmental stages, dauer neural network also exhibits its own form of variability (Figures 2C and 2D)(Witvliet *et al*., 2021). As the first distinctive characteristic of dauer neural network, the enhanced clustering and augmented triangular motifs (Figure 6B-6D), can allow faster signal transduction by making shortcuts, can enhance the robustness in the signal processing by providing an alternative path, or can amplify the signal by making a cyclic, recurrent connection (Peron et al., 2020). As the nictation seems to be initiated in the nose, our nerve ring connectome is potentially useful in elucidating the mechanism of nictation.

As a second distinctive characteristic, output connectivity of sensory neurons is dauer-specifically rewired compared to adults (Figures 6E-6G). While sensory connectivity is developmentally dynamic, strengthening or weakening existing connections, in the normal reproductive stages (Witvliet *et al*., 2021), dauer sensory neurons send signals to different target neurons (Figures 6E-6G), which could explain the differences in chemosensation. While dauers still show normal chemosensory responses to many chemical attractants as other reproductive stages, dauers also exhibit specificity (Cassada and Russell, 1975; Ward, 1973). For example, it has been reported *C*. *elegans* in the dauer stage respond differently to Na^+^ and CO_2_ (Altun and Hall, 2009; Gaglia and Kenyon, 2009; Golden and Riddle, 1984a; b; Hallem *et al*., 2011; Lee *et al*., 2011). These qualitatively different network structures may support the survival strategies of the dauer.

Unlike sensory and motor neurons, dauer interneurons show similar variability in connectivity to variability within different adult datasets (Figures 6E-6G). Wiring of interneurons are known to be developmentally stable, serving as the core central processor (Witvliet *et al*., 2021). Our results imply stability of interneuron connectivity could be conserved in the dauer stage as well, sharing the same central processor.

However, our classification of interneurons (Cook *et al*., 2019) are different from interneurons defined in (Witvliet *et al*., 2021) as it also includes modulatory neurons, which have many variable connections (Witvliet *et al*., 2021). In fact, connection similarity between adult-1 and adult-2 neurons were significantly greater than CS with dauer neurons (Figure S5E), contradicting the above hypothesis. For fair comparison, we have rerun the analysis without the modulatory neurons and discovered CS among datasets do not show significant difference (Figure S6), maintaining the hypothesis that the stability of interneuron connectivity is conserved in dauer.

In summary, as the last piece of the nematode connectome, we have reconstructed the first dauer nerve ring connectome from EM images and have shown that dauer connectome is quantitatively similar to and qualitatively distinct from the adult connectome. Our dauer connectome resource will open up many possibilities to study how the neurons function differently due to differences in the neural circuit, and vice versa.

### Limitations

As we have reconstructed only one dauer connectome, there exists a possibility that the features described in this study could be due to individual variability. We have conducted fluorescence imaging to validate certain features (e.g. IL2 morphological changes) also exist in other dauer animals but we could not validate all features. Gap junctions, another important component in the neural circuit, are not included in this study as it is difficult to identify gap junctions in our EM image dataset due to staining and fixation methods. Still, we have attempted to identify gap junctions that are obvious and the data is available along with the analysis code. Despite the above caveats, it is still the only publicly available dauer connectome dataset so we hope this resource could help the community in stage-wise comparative study of *C*. *elegans*.

## Acknowledgments

We thank Steven J. Cook and Oliver Hobert for reagents and valuable advice. We also thank Scott W. Emmons and Hannes E. Bülow for discussions. C. elegans strains were kindly provided by the Caenorhabditis genetic center (USA), Yuichi Iino. We thank Gwanho Ko and undergraduate students for their help with cell segmentation. This research was funded by the Samsung Science and Technology Foundation (Project Number SSTF-BA1501-52). H.Yim and D.T.C. were supported by the BK21 program. J.A.B. acknowledges support by the National Research Foundation of Korea grant (2019R1A6A1A10073437) funded by the Korean government (MEST).

## Author Contributions

K.N. generated EM dataset after sample preparation by H.K. B.S. stitched and aligned the images. H.K. segmented the EM images and H.Yim proofread them. J.A.B. trained and applied nets for synapses using ground truth generated by H.Yim, D.T.C. H.Yim analyzed morphology and connectivity differences in dauer with help from D.T.C and J.A.B. D.T.C analyzed network characteristics with input from J.A.B. and H.Yim. S.A. and H.Yim performed confocal experiments. M.C. conducted optogenetics experiments. M.C. and H. Yang performed nictation experiments. D.H. supervised image collection. H.Yim, D.T.C., J.A.B., D.H., J.S.K., and J.L. wrote the paper. J.L. and J.S.K. managed the project.

## STAR⍰Methods

### Experimental model and subject details

*C. elegans* were cultured and handled at 20°C according to standard methods (Brenner, 1974). *C. elegans* strains used in this study are as follows: N2 Bristol. JN2113: Is2113[gcy-21p::mCaspase1, tax4p::nls::YC2.60, lin44p::GFP]. These strains were obtained from the Caenorhabditis Genetics Center (CGC). JN2114 (Is[gcy-21p::mCaspase, gcy-21p::mCherry, lin44p::GFP]) was provided by Iino Yuichi’s lab. For TEM fixation, *C*. *elegans* Bristol strain N2 were used.

### Dauer induction

For TEM fixation, seven to eight young adult larvae were transferred to synthetic pheromone plates containing agar (10⍰g/L), agarose (7⍰g/L), NaCl (2⍰g/L), KH2PO4 (3⍰g/L), K2HPO4 (0.5⍰g/L), cholesterol (8⍰mg/L), and synthetic pheromone-ascaroside 1, 2, 3 (2⍰mg/L each) (Butcher et al., 2007; Jeong et al., 2005) seeded with Escherichia coli OP50 at 25⍰°C for dauer induction (Lee et al., 2015). After 4 days, dauers were induced and morphologically identified by their radially constricted bodies and sizes. For nictation assay, only synthetic pheromone-ascaroside 2 (3⍰mg/L each) were used for dauer induction.

### Fixation and staining for dauer larvae

Given the thicker cuticle on the dauer, we utilized an enhanced “OTO” protocol (Seligman et al., 1966) to conduct multiple rounds of osmium tetroxide exposure following a high-pressure freeze fixation to add extra layers of osmium stain prior to finishing a more regular counterstain with uranyl acetate and lead aspartate. The cryofixation by high-pressure freeze and freeze-substitution (HPF-FS) better preserves membrane quality, while the OTO adds to the final contrast compared to a single exposure to osmium.

Live dauers were immediately transferred into a bacterial slurry in a type A hat, matched to a sapphire disk to close the volume, backed by a type B hat (flat side towards sapphire disk) and mounted into the specimen holder of a Bal-tec HPM010 High Pressure Freezer. The samples were frozen under high pressure and moved to the first fixative, 5% glutaraldehyde, 0.1% tannic acid in acetone, at liquid nitrogen temperature. Multiple samples from the same run could be fixed in the same tube to conserve space in the freeze substitution device, and assure that portions of samples are well-frozen.

Frozen samples were moved into a Leica freeze substitution device at a holding temperature of - 90°C and held at that temperature for 64 hours, then warmed to −60°C in 5°C steps and held for another 48 hours. During this time, samples were washed several times in dry cold 98% acetone, then shifted to the second fixative, 2% osmium tetroxide, 0.1% uranyl acetate, 2% double-distilled (DD) water in acetone. After 48 hours, temperature was then raised slowly to −30°C in 5°C steps and held at −30°C for 16 hours. Then, it was immediately followed by four washes in 98% cold acetone for 15 minutes each. Samples were removed from the freeze substitution unit and put onto ice, then transferred to 0.2% thiocarbohydrazide in acetone for one hour at room temperature. After four 15-minute washes in 98% acetone, the samples were moved to the third fixative, 2% osmium tetroxide, 0.1% uranyl acetate, 2% DD water in acetone on ice for one hour. After three 10-minute washes in 98% acetone at room temperature, the samples were stained in 1.5 ml of filtered 2% lead aspartate in acetone for 15 minutes at room temperature. The samples were transferred to microporous specimen capsule (EMS, cat # 70187-21) for further handling of multiple worms without sample losses.

The samples were washed four times in 100% acetone for 15 minutes each at room temperature. Stepwise infiltration with resin:acetone dilutions (1p:2p and 2p:1p) was performed. Individual animals were mounted on Lab-Tek chamber slides with cover glass slide (EMS, cat # 70411) as molds. Then, cover glass was removed and the samples were cured in a 60°C oven for 48 hours, which makes it possible to cut out single animals from a thin layer of resin and Aclar to remount for thin sectioning.

### Electron microscopy image acquisition for dauer larvae

RMC PowerTome XL Ultramicrotome (Boeckeler Instruments, Inc.) and a 45-degree Ultra Diamond Knife (Electron Microscopy Sciences, cat # 40-US) were used for sectioning at a speed of 1 mm/s to cut 50 nm serial sections. The sections were collected onto Formvar-coated slot grids by hand (Hall and Rice, 2015).The sections were post-stained, first with 2% uranyl acetate for 20 minutes, and then a 1:5 dilation of Reynold’s lead citrate for 5 minutes.

Transmission electron microscopy imaging was conducted on a JEOL JEM-1400Plus microscope, equipped with a Gatan Orius SC1000B bottom mount digital camera. Using Serial EM software, we collected about 70 individual images per thin section.

### Stitching and alignment

Stitching and alignment for dauer EM images were done using TrakEM2 software (Cardona et al., 2012). First of all, image patches were normalized to have similar contrast and brightness. Scale-invariant feature transform (SIFT; (Lowe, 1999)) was applied to identify local features in the overlapping area of the adjacent image patches. Based on identified features, optimal affine transformation was computed, which minimizes least squares error, then applied to stitch image patches.

To align the stitched image sections, the total image stack was divided into multiple chunks of image sections and each chunk was aligned separately to prevent large distortion. For each chunk, image features were manually selected from the first and last sections, then the affine transformation parameters that minimize least square error were computed. Transformations for intermediate sections were computed by interpolation according to the section’s distance from the first and last sections. Affine transformation with manual features was applied to align the chunks. Lastly, elastic alignment in TrakEM2 was applied for the final alignment of the image stack.

### Volumetric reconstruction and cell identification

All neurons, muscle cells, and glia in the EM volume were volumetrically segmented. Using VAST (Berger et al., 2018), annotators manually traced each neuron and colored the cross-section of neuron. The direction of left-right axis was determined with the asymmetric ventral nerve cord. The neurons were identified by anatomical features such as location of soma, neurite trajectory, and distinctive morphological traits using other published EM datasets as references (Brittin *et al*., 2021; Cook *et al*., 2023; Kim et al., 2019; White *et al*., 1986; Witvliet *et al*., 2021). Neuronal types were defined by classification used in (Cook *et al*., 2019).

### Chemical synapse detection and synaptic partner assignment

Two areas of the volume have been selected from the EM volume and 10 sections from each area have been chosen to be annotated for the ground truth labels. Human annotators painted presynaptic active zones, which are darkly stained pixels near the membrane, using VAST (Berger *et al*., 2018) and the results have been reviewed by another annotator. Ground truth labels covering an area of 9053 μm^2^ have been generated. The labels have been divided into train (7340 μm ^2^), validation (874 μm^2^), and test (839 μm^2^) datasets.

The network used for the chemical synapse detection was adopted from 2D symmetric U-Net architecture (Olaf Ronneberger, 2015). The architecture was composed of five layers with a number of feature maps, 16, 32, 64, 128, 256 from the top most layer to the bottom most layer, respectively. For each step in downsampling and upsampling layers, the architecture consisted of three 3 x 3 non-strided same convolutions. In the downsampling layers, max pooling was used to downsample by a factor of 2 in each layer. In the upsampling layers, a transposed convolution layer with nearest neighbor interpolation followed by 2 x 2 non-strided convolution was used to upsample by a factor of 2 in each layer. Skip connection was included in every layer which concatenates feature maps at the same level of the left-hand side to the output of the transposed convolution layer. Instance normalization (Ulyanov et al., 2020) and rectified linear unit (ReLU) was added after each 3 x 3 convolution. At the end of the symmetric network, 3 x 3 non-strided same convolution and sigmoid function was applied to produce the same-sized output image, where each pixel value represents probability of each pixel belonging to the presynaptic active zone. The network takes an input EM image that has patch size of 576 x 576 pixels in 2 x 2 nm^2^ resolution and outputs an image with size of 512 x 512 pixels in the same resolution. Note that the input image is 64 pixels larger in each dimension as 32 pixels from the edges have been cut off to avoid inaccurate predictions near the borders.

The network has been trained using an Adam optimizer (Diederik P. Kingma, 2014) and using binary cross entropy as a loss function. The learning rate has been kept constant at 0.0005. Horizontal and vertical flip and multiple of 90-degree rotation were applied on training samples as data augmentation. Also, brightness and contrast augmentation have been applied.

From the predicted probability map of active zones, we thresholded the image with a pixel threshold of 90 / 255 (35%) and generated binarized prediction images of active zones. To reconstruct active zones in 3D, we applied connected components with connectivity of 26. The errors in resulting reconstructions were corrected manually.

For each active zone, we ran Monte Carlo simulation of neurotransmitter diffusion (Witvliet *et al*., 2021) to identify potential synaptic partners. We assigned synapses sizes for each synaptic partner according to the proportion of particles that reached its segment.

### Generation and modification of transgene constructs

All constructs in this study were created using Gibson assembly cloning kit (E5510; New England Biolabs). DNA fragments were inserted into the GFP vector (pPD95.77) and mCherry vector (pPD117.01; modified to mCherry red fluorescence). For transcriptional fusion constructs, we inserted the promoter region of relevant genes into vectors. The indicated numbers are relative to the start codon ATG for each gene, respectively: *egas-4p::gfp*; *twk-3p::Chrimson::*SL2::*mCherry*. All details of transgene constructs are available upon request.

### Generation of transgenic animals

To generate transgenic animals, we microinjected DNA and plasmid constructs into the gonad of young adult hermaphrodite as a canonical method (Mello et al., 1991). We used following co-injection marker with indicated concentration for transgenesis to isolate transgenic progeny of microinjected hermaphrodite: *unc-122p::ds-red* (25 ng/ul; red fluorescent coelomocyte) and *myo-3p::mCherry* (50 ng/ul; red fluorescent muscle). Plasmid DNAs used for microinjection were extracted and purified using a plasmid purification kit. All details of transgenic animals are available upon request.

### Fluorescence microscopy

Images of worms were acquired using the confocal microscope (ZEISS LSM700; Carl Zeiss). For imaging, transgenic worms were paralyzed with 3 mM levamisole and mounted on 3% agar pads. To examine the morphological and synaptic changes of neurons, we visually inspected neurons by maximum intensity projection using ZEN software (Carl Zeiss).

### Contact matrices and contact area

In non-dauer datasets except for adult-1 dataset, which does not have segmentation images, segments in the region from where the nerve ring begins to the entry point of the ventral nerve cord (approximately 12∼13 μm length in L2 and L3) were used to compute contact matrices for fair comparison between different size image volumes. Reasonable background masks were obtained by manually annotating the dilated results of all segments. Each segment was dilated one at a time in pseudo-random order repeatedly until the background masks were fully covered. As non-dauer datasets do not have cell bodies and long amphid processes segmented, these segments were removed from the dauer segmentation images when computing contact matrix.

To measure the contact area between cell segments, contact direction (i.e., x-y, x-z, or y-z) was identified for each voxel in the contact region. For each contact voxel, the area of the square orthogonal to the contact direction was measured, and contact area was calculated as the sum of the areas for all contact voxels. For fair comparison of contact areas between datasets, contact areas have been normalized by the standard deviation of connection weights without the top 5th percentile to remove the bias due to the big outliers. Then, these contact areas and normalized contact areas have been assigned to corresponding locations in the 187 × 187 (neurons) common contact matrix structure for all datasets to create contact matrices.

### Connectivity matrices and connection weights

Connection weights between a connected pair of neurons were computed as the sum of all synapse sizes of synapses between pre- and postsynaptic neurons of the pair. For fair comparison of connection weights between datasets, connection weights have been normalized by the standard deviation of connection weights without the top 5th percentile to remove the bias due to the big outliers. Then, these connection weights and normalized connection weights have been assigned to corresponding locations in the 187 × 187 (neurons) common connectivity matrix structure for all datasets to create connectivity matrices.

### Wiring diagrams

Network diagrams were drawn using Cytoscape (Shannon et al., 2003). In the diagram, edges with normalized connection weights below the average of individual synapse weights (*w*=0.2728) were excluded. Also, connections between neurons and CEPsh cells and autapses, synapses to itself, have been excluded from the diagram. Connections are drawn as arrows, indicating the direction of connections, and the width of arrows represent the connection weights. In the figures with two wiring diagrams for comparison, the color of the edges indicates the changes in normalized weights in the dauer stage. Red and blue indicate increase and decrease in the dauer stage respectively. Orange indicates dauer-specific connection.

### Stage-conserved and stage-specific connections

Connections included in the intersection of all 11 datasets (Figure 2B) were assigned as stage-conserved connections. In each dataset, the connections that only exist in the dataset and cannot be found in any of the other 10 datasets were assigned as stage-specific connections.

### Nictation assay

First, we created a micro-dirt chip by pouring a 3.5% agar solution onto a PDMS mold (Lee *et al*., 2015). The solidified chip was then detached from the PDMS mold and dried for 90 min at 37°C. For individual nictation assays, individual dauer were observed for 1 min and measured nictating time. The nictation ratio was calculated from the ratio of nictating time and total observation time. Quiescent dauers were excluded from measurements. For population nictation assays, more than 30 dauer larvae were collected by a glass capillary tube using M9 isotonic buffer and mounted onto a freshly prepared micro-dirt chip. After 10–30 min, when dauers started to move, nictation was quantified as the fraction of nictating worms among moving dauers. Nictation assays were carried out at 25°C with a humidity of 30%. Assays were repeated at least four times for quantification and statistics. (Lee *et al*., 2015; Lee *et al*., 2011). For optogenetic experiments, young adult worms expressing Chrimson transgenes were transferred to either normal OP50 pheromone plates or OP50-retinal pheromone plates containing 1 mM all-trans-retinal (Sigma). The transgenic dauers derived from the pheromone plate were transferred to a micro-dirt chip and their behavior was recorded for approximately 1 minute under green light (510-560 nm) on a Leica M205 FA microscope. The behavior of a dauer from the OP50-retinal pheromone plate and a dauer from the normal OP50 pheromone plate was recorded alternately, and the nictation ratio was quantified from the recorded movies.

### Network properties

Total of 185 neurons, excluding CAN neurons as they have no connections, were used to compute network properties. Among 185 neurons, 71 were sensory neurons, 73 were interneurons, and 41 were motor neurons. The analysis was restricted to a network of neuronal cells, excluding muscle cells and end organs, as non-neuronal cells have inconsistencies between datasets.

Out- and in-degrees of a neuron was calculated by counting the number of postsynaptic partners and presynaptic partners, respectively. Out-degree for each connection type was calculated by counting the number of postsynaptic partners of in that subnetwork. Neurons without any connection in any of the dataset were neglected when computing out-degrees of different connection types.

Path length between a pair of neurons is computed by counting the minimum number of steps from the source neuron to the destination neuron.

Clustering coefficient (CC) measures the probability of the pair of neurons that are connected to a common neuron being connected to each other. CC was defined as

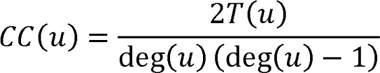

where *deg*(*u*) is the number of edges connected to the neuron regardless of their directions. *T*(*u*) indicates the number of triangles via node *u*, which occurs when the two neurons connected with neuron *u* are connected to each other. Nodes with less than two edges were excluded from the analysis as they have no chance of forming a triangle. Besides measuring the CC of the whole network, CC has been also measured in the sensory, interneuron, and motor subnetworks which only consist of connections within the same type of neurons.

### Number of common connections

Number of common connections measures the number of outgoing connections of a neuron that exist in both datasets being compared. The number of common connections has been measured separately depending on the type of presynaptic neurons and connections to all types of postsynaptic targets were taken into account. Neurons which do not have any outgoing connections in both of the datasets being compared were excluded from the analysis. In Figure 6E, the number of common connections for adult vs. adult was computed by averaging the values in three possible pairs (adult-1 vs. adult-2, adult-1 vs. adult-3, adult-2 vs. adult-3) and the number of common connections for dauer vs. adult was computed by averaging the values in three possible pairs (dauer vs. adult-1, vs. adult-2, vs. adult-3).

### Connection similarity

Connection similarity (CS) measures the amount of similarity between two connectivity vectors of a neuron. CS was defined as

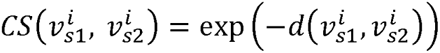

where 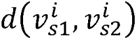 is a Euclidean distance between two connectivity vectors of neuron *i* (i.e., binary vector of outgoing connections) from stages *S*1 and *S*2. Exponential to the negative power is applied so vectors with greater distances can have lower similarity values and to force the range of the measure between 0 and 1. As the number of common connections, connection similarity has been measured separately depending on the type of presynaptic neurons. Neurons which do not have any outgoing connections in both of the datasets being compared were excluded from the analysis. In Figure 6F, connection similarity for adult vs. adult was computed by averaging the values in three possible pairs of adult datasets (adult-1 vs. adult-2, adult-1 vs. adult-3, adult-2 vs. adult-3) and connection similarity for dauer vs. adult was computed by averaging the values in three possible pairs of dauer and adult datasets (dauer vs. adult-1, vs. adult-2, vs. adult-3).

### Motif analysis

The whole network has been divided into three subnetworks: sensory, interneuron, and motor subnetworks. Then, the analysis has been done separately for each subnetwork, which only consists of connections within the neurons of that type. To compute the occurrences of three-cell motifs, all combinations of three cells were extracted from the binarized connectivity matrix. For each combination, it was assigned to one of 16 motif patterns depending on its connectivity. The proportion of triangular motifs was computed by dividing the sum of numbers of motifs that form triangles (motifs in black rectangle in Figure 6B) by the total number of three-cell combinations among 185 neurons. Same computation has been done for 1000 ER random networks, which have the same number of nodes and connection probability, for comparison.

### Statistics

Statistical tests were done by custom Python script using SciPy library. Unpaired t-tests were used to compare the results of behavior experiments (Figure 3C-3E). Data point for dauer was excluded when measuring correlation since it was difficult to decide the proper position for it. Two-sided Wilcoxon rank-sum tests were used to compare distributions of connection weights (Figure S3E) and network properties (Figure 6).

### Data and code availability

All data are available in the main text or the supplementary materials. Requests for resources should be directed and will be fulfilled by the corresponding author, Junho Lee.

## Supplementary Information

**Supplementary Table 1. Neuroglancer link and segmentation keys.**

Neuroglancer link for the visualization of reconstructed dauer data. 3D reconstructed cells can be visualized with 2D raw EM, cell segmentation, and synapse segmentation images. Segmentation key table includes segment ids of the reconstructed cells in the neuroglancer.

**Supplementary Table 2. Dauer connectivity matrix.**

Connectivity matrix of dauer connectome with raw connection weights (nm3) and normalized connection weights (see Methods). The rows represent presynaptic cells and the columns represent postsynaptic cells. Connectivity matrices of other datasets used in the study are also included as a reference.

**Supplementary Table 3. Dauer-specific, dauer-loss, stage-conserved connection lists.**

List of dauer-specific, dauer-loss, and stage-conserved connection lists. The normalized weights of each connection in dauer and adult-2 are shown, which have been used to draw comparative wiring diagrams.

**Supplementary Table 4. Dauer contact matrix.**

Contact matrix of dauer connectome with raw contact area (nm2) and normalized contact area (see Methods). The contact matrix is a symmetric matrix as there is no distinction between presynaptic and postsynaptic cells. Contact matrices of other datasets used in the study are also included as a reference.

**Supplementary Item 1. Dauer wiring diagrams.**

Wiring diagrams of all neurons. The colors of the nodes represent the cell types (pink: sensory neuron, orange: interneuron, blue: motor neuron, green: body wall muscle). The width of the arrows indicates the relative connection weights. Dauer-specific connections are represented with orange. Connections which the weight increased and decreased in dauer are represented with red and blue arrows, respectively.

---

**Figure S1.**
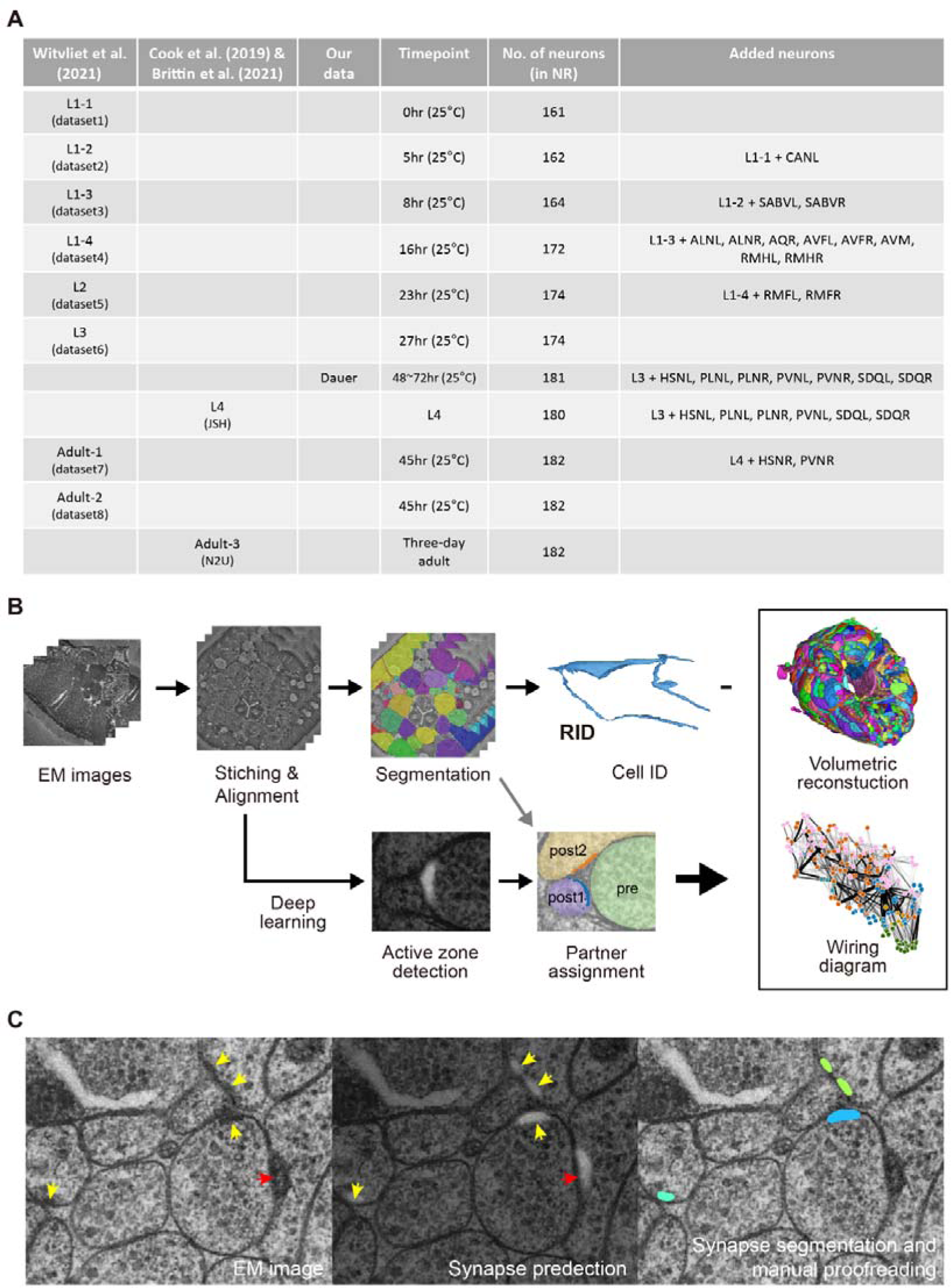
Reconstruction pipeline and comparison with other datasets, related to Figure 1. (A) EM connectome datasets in different developmental stages used for comparison. The number of neurons present in the nerve ring increases as the worm matures. PVNR neuron already exists in the dauer stage while it does not exist until L4 stage and appears in the adult stage. (B) Reconstruction pipeline of dauer connectome: alignment of EM images, cell reconstruction and identification, and synapse detection. Electron microscopy (EM) image tiles were stitched and the stitched sections were aligned. From the aligned image stack, annotators manually traced and colored the area inside the membrane for each cell. Classes of reconstructed cells were identified according to morphological features. Chemical synapses were identified by detecting active zones in the EM images using convolutional neural network (CNN). For each active zone, the synaptic partners were assigned, giving a complete connectivity graph of reconstructed neurons. (C) Synapse detection in sample EM image cutout. Probability of pixels being in the darkly stained presynaptic active zones (yellow arrow) in the EM image (left) are predicted using CNN (middle, white indicates higher probability). Then for each predicted active zone, the probability values were thresholded and grouped using connected components to segment individual active zones (right, labeled with different colors). Lastly, manual proofreading has been done to remove falsely predicted active zones (red arrow) or falsely assigned active zones.

**Figure S2.**
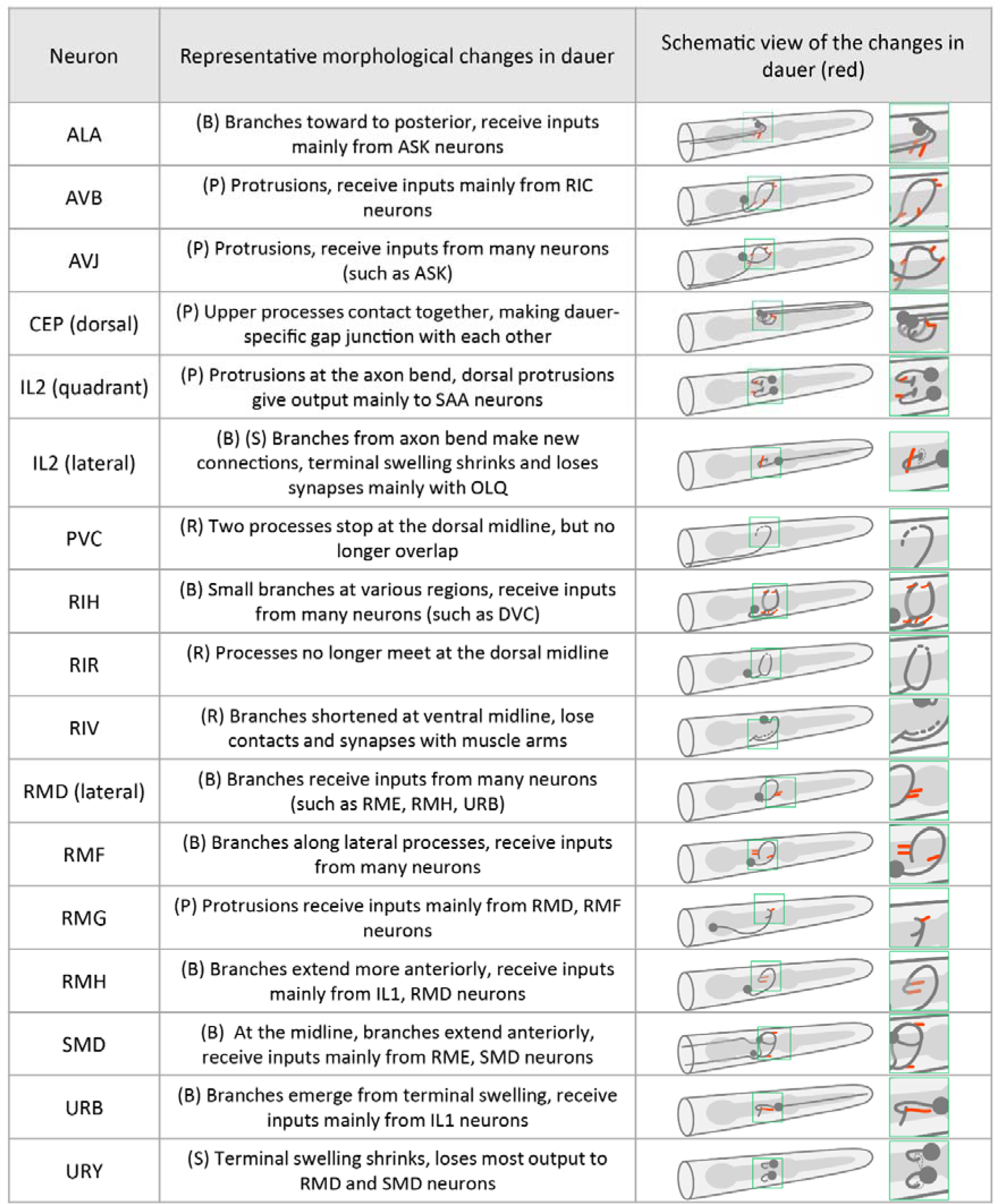
Morphological changes of neurons in the dauer stage, related to Figure 1. Summary of representative morphological changes of neurons in the dauer stage with schematic. Morphological changes are classified into 4 types: branching (B), small protrusion (P), retraction (R), and shrinkage (S).

**Figure S3.**
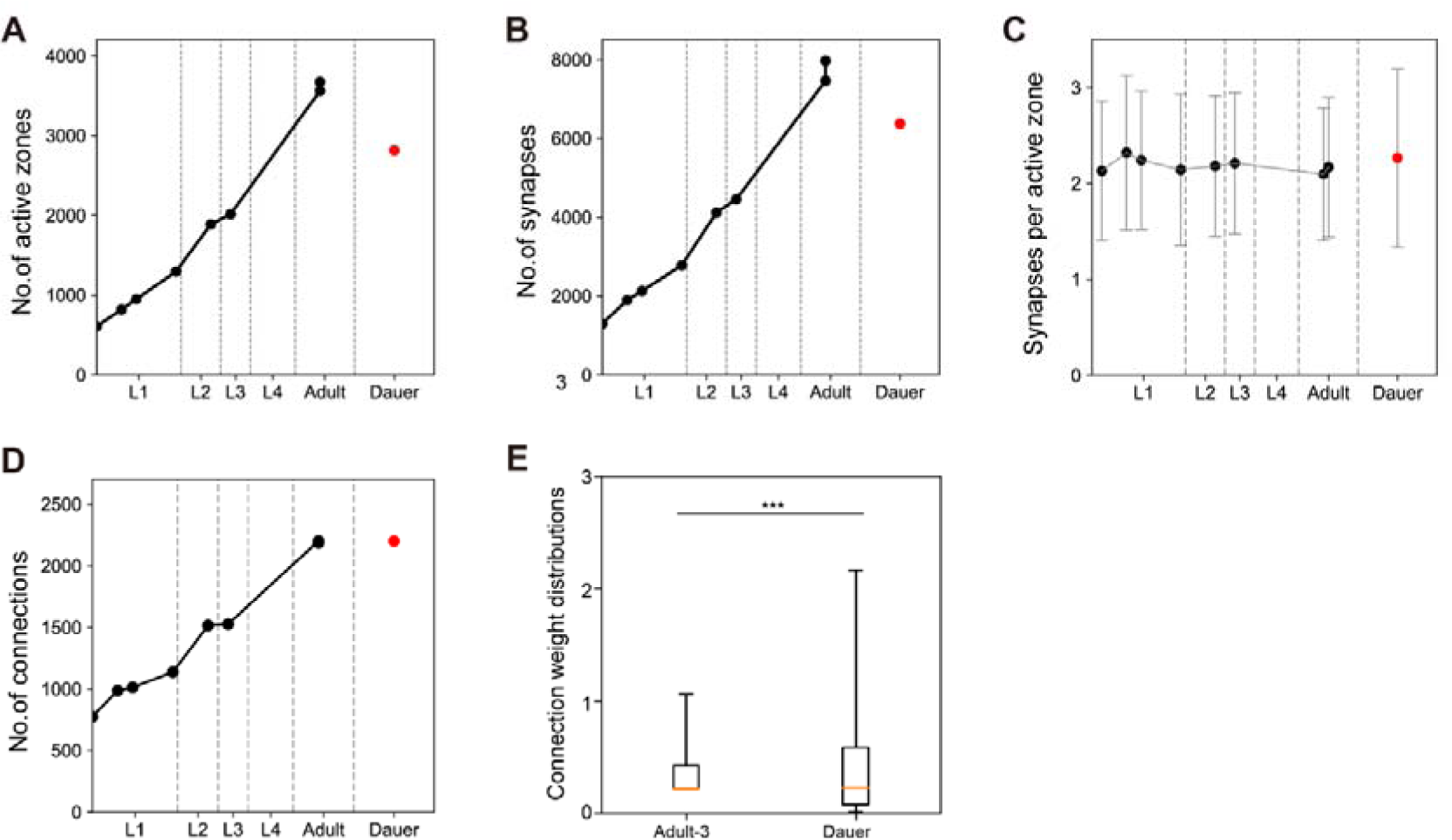
Connection properties across development, related to Figure 2. (A) Total number of active zones across development. L4 and adult-3 datasets are excluded as they do not provide synapse information. (B) Total number of synapses across development. (C) Average number of postsynaptic partners per active zone is consistent across development including the dauer stage (mean±SD). (D) Total number of connections including neuromuscular junctions across development. (E) Normalized connection weight distribution of stage-specific connections in dauer dataset and adult-3 dataset (Orange line: median, box: interquartile range, whiskers: 5th and 95th percentiles; Wilcoxon rank-sum test; *p*=5.02E-08).

**Figure S4.**
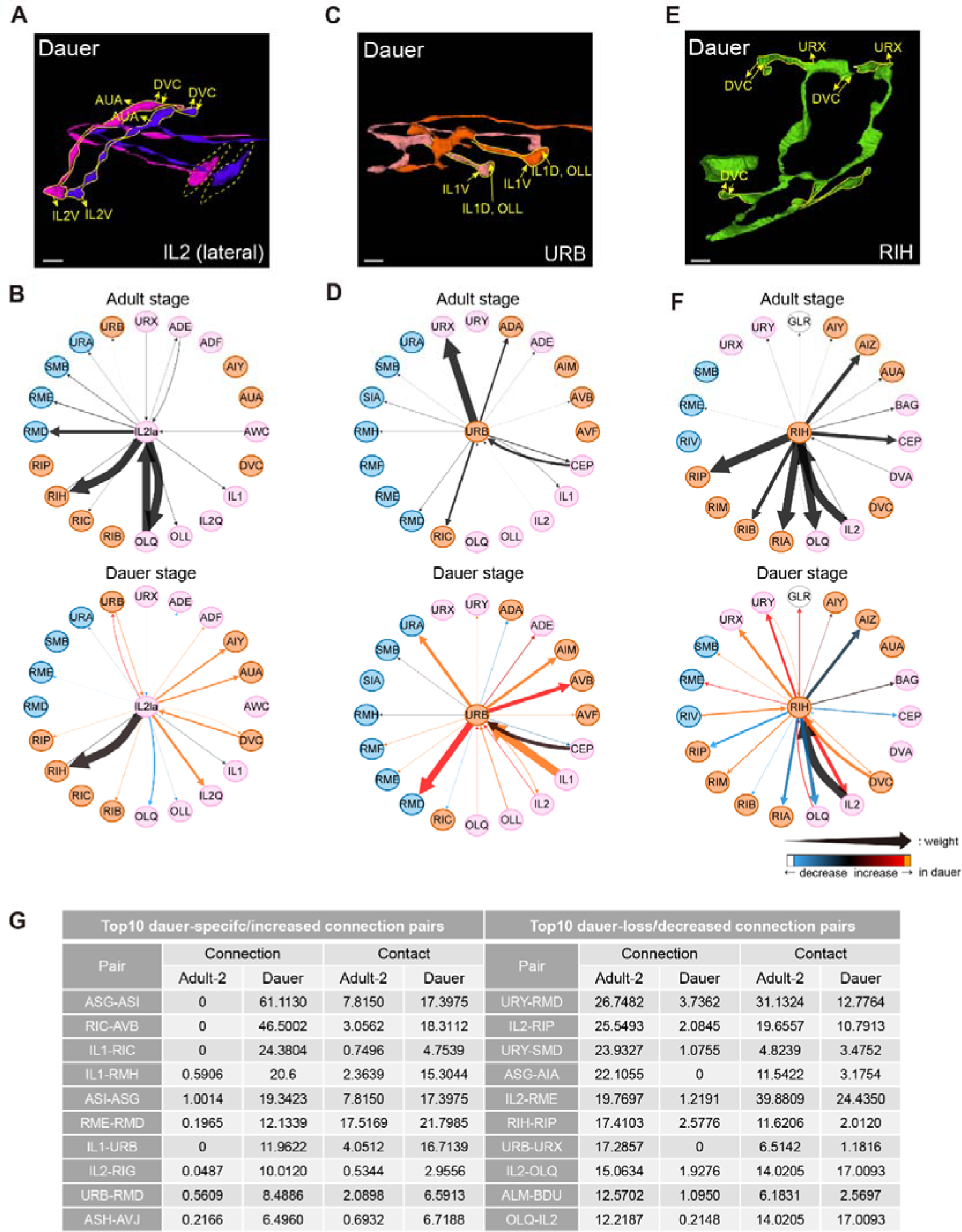
Morphological changes are accompanied by new synaptic inputs and outputs, related to Figure 3. (A) 3D view of IL2 lateral neurons in dauer with new synaptic inputs and outputs (yellow arrow) in the region where morphological changes occurred (solid yellow line). (B) Wiring diagram of IL2 lateral neurons in adult (top) and dauer stages (bottom). Width and color of the arrow follow the same rule from Figure 2D. (C) Same with (A) for URB neurons. (D) Same with (B) for URB neurons. (E) Same with (A) for RIH neurons. (F) Same with (B) for RIH neurons. (G) Normalized connection weights and contact areas for 10 pairs of neurons with greatest increase in connection weights (left) and greatest decrease in connection weights (right) in dauer relative to adult-2 dataset. (A, C, E) Scale bars: 1 μm.

**Figure S5.**
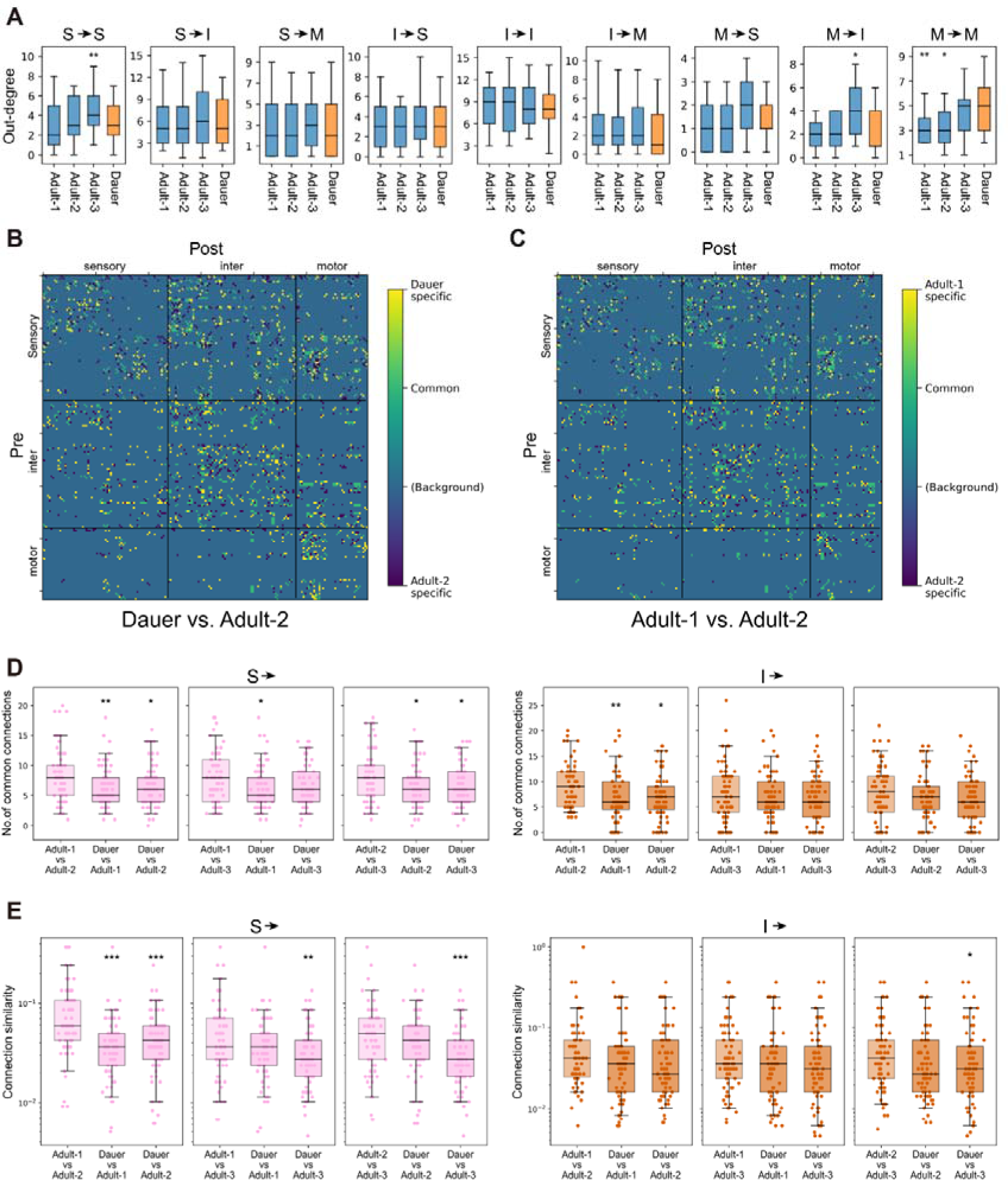
Comparison of connectivity in dauer and adult networks, related to Figure 6. (A) Expanded version of Figure 6A (*n*=185; Wilcoxon rank-sum test; \**p*<0.05). (B and C) Connectivity difference matrices between dauer and adult-2 (B) and that between adult-1 and adult-2 (C). Connections that are shared between datasets are marked as green and stage-specific connections are marked as yellow or dark blue. Dauer-specific connections (yellow in (B)) are dominant in the motor subnetwork (lower right corner). (D) Expanded version of Figure 6E for sensory (pink) and inter- (orange) neurons (Wilcoxon rank-sum test; \**p*<0.05, \*\**p*<0.01). (E) Expanded version of Figure 6F for sensory (pink) and inter- (orange) neurons (Wilcoxon rank-sum test; \**p*<0.05, \*\**p*<0.01, \*\*\**p*<0.001). (A, D, E) Black line: median, box: interquartile range, whiskers: 5th and 95th percentiles.

**Figure S6.**
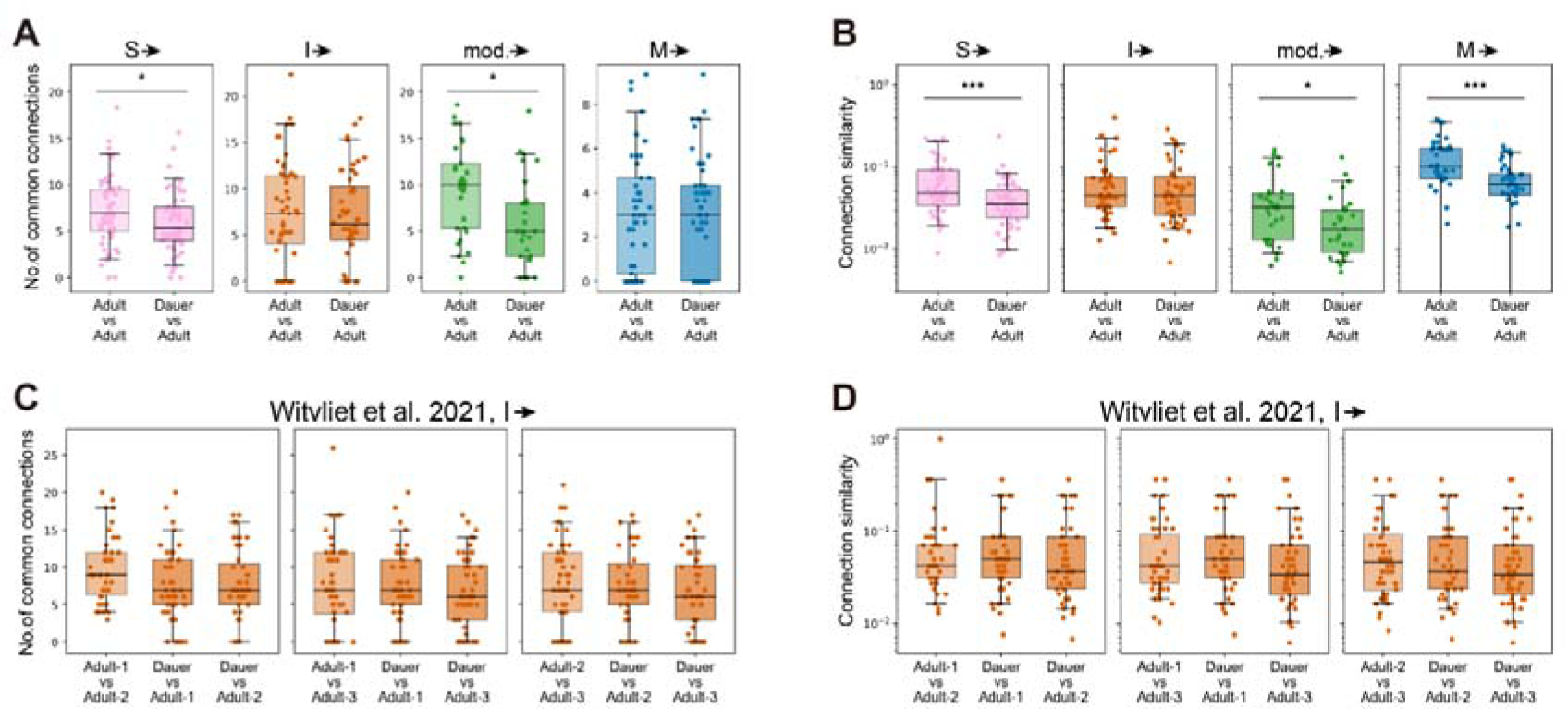
Comparison of connectivity in dauer and adult networks using type classification from Witvliet et al. 2021, related to Figure 6. (A) Number of common output connections from sensory (pink), inter- (orange), modulatory (green), motor (blue) neurons, defined in Witvliet et al. 2021, between adults (left) and that between dauer and adults (right; Wilcoxon rank-sum test; \**p*<0.05). (B) Connection similarity of output connections from sensory (pink), inter- (orange), motor (blue) neurons, defined in Witvliet et al. 2021, between adults (left) and that between dauer and adults (right; Wilcoxon rank-sum test; \**p*<0.05, \*\*\**p*<0.001). (C) Expanded version of Figure S6A for interneurons defined in Witvliet et al. 2021. (D) Expanded version of Figure S6B for interneurons defined in Witvliet et al. 2021. (A-D) Black line: median, box: interquartile range, whiskers: 5th and 95th percentiles.

